# Numerical simulations of different models describing cerebrospinal fluid dynamics

**DOI:** 10.1101/573345

**Authors:** Licia Romagnoli

## Abstract

The aim of this paper is to present an extensive overview of numerical simulations aimed at confirming and completing the theoretical results obtained in the analysis of some cerebrospinal fluid dynamics models which are treated from a purely mathematical view point. The present study is designed to support the first attempts in the approach to these physiological models from a more theoretical standpoint since their investigation in literature only concerns the modelization and the clinical feedback.

## 1. INTRODUCTION

The cerebrospinal fluid (CSF) perfuses the cerebral ventricles and the cranial and spinal subarachnoid spaces (SAS). An anatomical overview of spaces filled with cerebrospinal fluid is shown in Fig. 1. This physiological body fluid is not static, but shows a pulsating movement within the ventricular system and between the cranial vault and spinal compartments, overlapped by the bulk flow affected by the fresh production and by final reabsorption in venous systems. In order to guarantee the normal brain function, the pulsatile pattern is fundamental but several diseases are able to destabilize the complex intracranial equilibrium by affecting the CSF flow dynamics.

**Figure 1:**
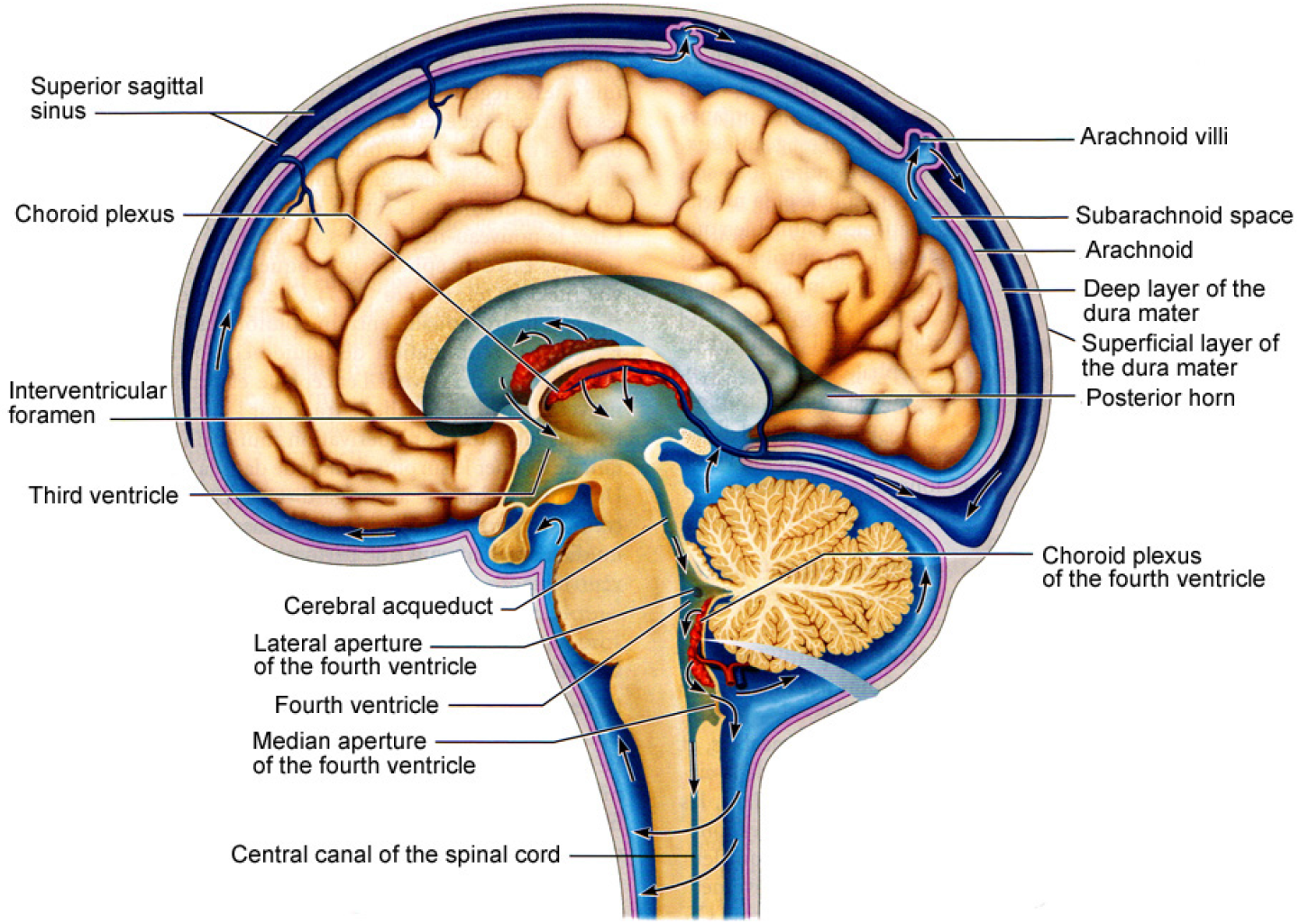
Cerebrospinal fluid circulation.

**Figure 2:**
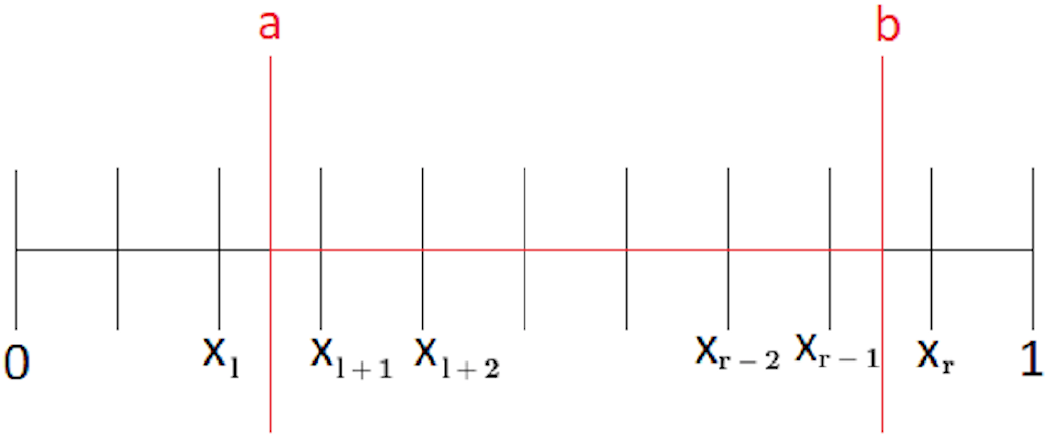
Discretization of the domain.

Recent efforts in scientific and clinical communities are aimed precisely at quantifying the critical parameters relating to normal intracranial dynamics and at identifying the deviations characterized by the diseases. The intracranial pressure (ICP) is extremely relevant but the only ICP measurement technique available at the moment is in vivo. Furthermore, CSF flow velocities, the motion of the brain and the deformations which take place inside the skull and the spinal compartments can be achieved without applying invasive techniques.

In vivo medical imaging provides an impressive overview in the intracranial and spinal dynamics. Image data are essential in order to accurately define anatomical spaces of the patient, to determine blood and cerebrospinal flow, and to track the biodistribution of pharmacological agents transported by the pulsatile CSF. Since in vivo imaging often guarantees only measurements in a single localized point, researchers and clinical operators are forced to hypothesize about the physiological and biomechanical interactions between the central nervous system (CNS) compartments. Nevertheless, speculations without quantitative proof have so far not consolidated our knowledge about brain disease enough to improve the treatment of subjects affected by diseases.

Quantitative models are fundamental to provide difficult predictions in order to support new intuitions about the fluid dynamics of blood and cerebrospinal fluid in the central nervous system. In this scenario, the important role of mathematical models is evident since they are able to achieve a better interpretation about in vivo data detected at multiple length scales and anatomical positions inside the intracranial pattern.

The purpose of this paper is to present an overview of numerical simulations aimed at confirming and completing the theoretical results obtained in [8] and [9], where the cerebrospinal fluid (CSF) dynamics models analyzed are mainly treated from a mathematical point of view. In the mentioned papers the authors study CSF models introduced by Linninger et al. ([21]) and Marmarou et al. ([25], [27]), and prove first local existence and uniqueness of solutions for the systems of equations which rule the analyzed dynamics, then they investigate the global solutions by focusing on the nonlinear equation describing the CSF flow velocity. In particular, it is proved that these solutions exist and are unique under proper restriction conditions on the flow velocity and intracranial pressure initial data.

We will carry out the numerical simulations based on three major topics: a good approximation of the equations, a numerical method which is not expensive in terms of computational costs, boundary and initial conditions which fulfill the mathematical analysis of the cerebrospinal fluid dynamics models treated in [8] and in [9]. In order to assess the usefulness of the theoretical analysis developed in the aforementioned papers, we discuss the numerical results from a qualitative standpoint and when available, we compare them with quantitative information derived from literature or experimental data, even though some assumptions are a bit far from the real physiology of the cerebrospinal fluid.

Section 2 begins with a current view of the intracranial dynamics connecting the cerebrospinal fluid to the vascular pulsation. In Section 3 we move from a description of the parameters and the quantities that are involved in all models we will treat numerically, and the range of values they are allowed to assume throughout the particular CSF dynamics process described by the models. Moreover, we will introduce the three compartmental CSF models (**Model A, A1, A2**) that represent the core of the mathematical analysis of [8] and [9] for which we want to perform proper numerical simulations. Every model will be supplemented by suitable initial and boundary conditions that, wherever possible, will comply with the clinical data.

Then, in Section 3, we will point out the goal of the present paper.

Furthermore, in Section 4, we proceed by defining the numerical scheme for the three models and we carry out the numerical simulations in two different cases: first, we fix initial data which satisfy the conditions required by the global existence theorems proved in [8] and in [9], then, we choose initial data that violate them.

Finally, we conclude with some fundamental considerations in Section 5.

## 2. Hints on physiological intracranial dynamics

### 2.1 Cerebrospinal fluid motion in the cranial vault

The cerebrospinal fluid is a clear fluid with density and viscosity similar to those of water. It surrounds the brain and spinal cord and its protein content is lower than that of blood plasma ([30]). CSF perfuses the ventricular system and the cranial and spinal SAS. Moreover, within the central nervous system CSF is not stagnant but it pulsates through the lateral ventricles. Thanks to the imaging studies, researchers have confirmed that the cardiac cycle prescribes its pulsatile behavior on the CSF ([11]). During the systole CSF flows from cranial to the spinal SAS, with flow inversion from the spinal SAS towards the cranium in diastole. It has been observed also a respiratory influence on CSF oscillations in the aqueduct ([1], [10], [15], [31], [34], [35]).

Furthermore, it has been detected a small volumetric bulk component as well as marked fluctuations in the flow without a net flux. It is believed that the new CSF is secreted through the epithelium of the choroid plexus (see Fig. 1). Figure 1 shows how the lateral ventricles communicate with the third ventricle, and a thin tubular canal, the Sylvius aqueduct, links the third to the fourth ventricle. The flow in the aqueduct performs a strong pulsatile behavior ([32]). The foramina of Magendie and Luschka originate in the fourth ventricle and extend into the cerebral SAS at the prepontine area. Since the the human central nervous system cannot claim a classical lymphatic system, the clearance of CSF is different from the peripheral extracellular drainage of fluids. The most reasonable hypothesis is that CSF is reabsorbed into the venous system through the arachnoid villi, granulations of the arachnoid membrane in the superior sagittal sinus, which lies at the top of the head (see Fig. 1) or alternatively through nerve pathways in the extracranial lymphatic system.

Experiments performed on different animal species propose that lymphatic drainage is significant in rodents ([2], [16]) and dogs ([19], [24]). The most recent results indicate the existence of a meningeal lymphatic network in mice ([23]). Nevertheless, the extension of lymphatic drainage in humans is still debated ([14], [33]). The alternative theories of CSF production and reabsorption cast doubt on this traditional view ([4], [3], [18], [17], [28]). Klarica et al. [18], endorsed by dilution experiments in cats, suggested the theory that the production and reabsorption of CSF takes place throughout the parenchyma without a clear bulk component by means of choroidal production or clearance via the arachnoid villi.

### 2.2 Cerebrospinal fluid flow and cerebral vasculature

The bulk of the cerebrospinal fluid obtained by new production and reabsorption is small compared to its pulsatile component. Researchers hypothesize that the oscillations of the pulsatile CSF are driven by systolic vessels dilatation followed by diastolic contraction.

#### 2.2.1 Cerebral force interactions in intracranial pattern

In a normal cardiac cycle, approximately 750 ml of blood are pumped into the head per minute. The increase in systolic blood pressure inflates arterial blood vessels, therefore it is possible to detect an increase of the cerebral blood volume during systole. Since the cranial vault is enclosed in a rigid bone ossification in adults, the vascular expansion of the main arteries that pass through the spaces filled with CSF enables the displacement of CSF. MR imaging shows that the total volume of cerebral blood increases and decreases in every cardiac cycle of about 1 – 2 ml, the same volumetric quantity as there is the exchange of CSF between the cranial and spinal SAS ([12]).

Thanks to an MRI technique developed by Zhu et al. ([36]), it has been quantified the ventricle movement of the wall, which drives the flow of the cerebrospinal fluid into the ventricular system. Two case scenarios could be identified: dilatation of the vascular volume could be transmitted from the cortical surface through brain tissue, whose compression causes the contraction of the ventricular space; otherwise, ventricular wall motion could derive from inside the ventricles by systolic expansion of the choroidal arteries, due to the ventricular walls pulsation against the periventricular ependymal layer. Ventricular dilatation due to the expansion of the choroid plexi was hypothesized in the theoretical model of Linninger et al. ([21]) and measured with cine PC MRI ([36]).

As well as the arterial expansion, it has been hypothesized a compression of the venous system under high pressure, which would affect the cross-sectional area by deformation, by reduction of the venous blood lumen or even by collapse of venous tree sections. Greitz et al. ([13]) proposed that the venous system is compressible, especially in patients with high ICP. As a result, high intracranial pressures compress the venous bed mainly in the superior sagittal sinuses. The venous lumen decreases and generates feedback that reduces blood flow due to the resistance increment.

## 3. Preliminary considerations

One of the biggest challenges in the numerical resolution of the lumped models analyzed in [8] and [9] is setting up the many constants and parameters that characterize the problem and that affect the solution both quantitatively and, sometimes, qualitatively. In fact, in the literature, the experimental or theoretically predicted range for the values of many of these parameters is often quite large.

The description and the representation of the human brain anatomical structure need a detailed data set, even if the approach based on the first principle of fluid mechanics requires only a small number of physical and physiological parameters. The physical constants and the quantities involved in the CSF models we will treat numerically are defined in Table 1, while the values of constants including the CSF viscosity (*μ*), density (*ρ*) as well as spring elasticity (*κ*) and brain dampening 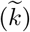 are listed in Table 2 ([22], [26]). In addition, a set of boundary conditions needs to be specified.

**Table 1:** Table of symbols and quantities involved in CSF models.

| NOMENCLATURE |  |  |
| --- | --- | --- |
| $\rho$ | fluid density | [kg/m<sup>3</sup>] |
| $r$ | radius of the foramina and aqueduct | [m] |
| $\mu$ | fluid viscosity | [Pa s] |
| $\beta = 8\mu/r^2$ | Poiseuille friction term | [N/m<sup>3</sup>] |
| $\delta$ | tissue width | [m] |
| $\tilde{h}$ | height of the ventricular or subarachnoid section | [m] |
| $Q_p$ | CSF production rate in the choroid plexus | [m<sup>3</sup>/s] |
| $R$ | resistance to CSF absorption | [Pa s/m<sup>3</sup>] |
| $\bar{\alpha}$ | amplitude of choroid expansion | [m] |
| $\omega$ | heart rate frequency | [rad/s] |
| $K$ | mathematical constant | [ml] |
| $\kappa$ | tissue elasticity constant | [N/m] |
| $\tilde{k}$ | tissue compliance | [N s/m] |
| $\tilde{P}$ | pressure of brain parenchyma | [N/m<sup>2</sup>] |
| $L$ | length of a single CSF compartment | [m] |
| $z$ | axial coordinate | |
| $A(t, z)$ | cross section of the ventricular or SAS | [m<sup>2</sup>] |
| $\eta(t, z)$ | tissue displacement in a section | [m] |
| $Q_a(t, z)$ | CSF absorption rate | [ml/min] |
| $u(t, z)$ | axial CSF flow velocity | [m/s] |
| $P(t, z)$ | CSF pressure in ventricles and SAS (ICP) | [N/m<sup>2</sup>] |

**Table 2:** Tissue and fluid properties.

|  | Value |
| --- | --- |
| $\rho$ | 1004 – 1007 kg/m<sup>3</sup> |
| $\delta$ | $\sim 5 \times 10^{-4}$ m |
| $\mu$ | 10<sup>-3</sup> Pa s |
| $\kappa$ | 8 N/m |
| $\tilde{k}$ | $0,35 \times 10^{-3}$ N s/m |

The bulk production of CSF takes place in the choroid plexus and in the neural tissue, in particular the parenchyma. The diffuse production of CSF flow throughout the parenchyma is accounted for as a source term, *Q_p_*. The bulk CSF production is implemented as input flux at the choroid plexus and has been chosen to correspond to the daily physiological production rate of 0.5 liters.

Recently, it has been shown (see [36]) that the expansion of the vascular bed in the systole leads to compression of the lateral ventricle as well as enlargement of the choroid plexus resulting in CSF flow out of the ventricles. For this study, the action of the vascular expansion bed is accounted for via the boundary condition for the choroid plexus as given by the following

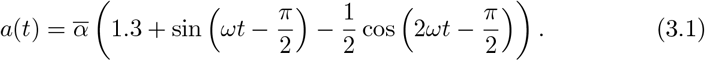

Thus, the choroid boundary condition accounts for the constant CSF production as well as the pulsatile flow of CSF due to expansion of the parenchyma as well as choroid plexus in the systole (see [20], [21]). The frequency of the pulsatile motion is set to 1 Hz approximating the normal cardiac cycle.

Most scientists believe that the majority of reabsorption of the CSF is into the granulation of the sagittal sinuses. Accordingly, re-absorption of the fluid takes place at the top of the brain geometry. The re-absorption is assumed to be proportional to the pressure difference between the ICP in the SAS and the venous pressure inside the sagittal sinus (see [25]). This relation is expressed mathematically by

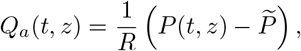

where the resistance *R* has physiological values of the order of 10 ml/(mmHg min).

In what follows, we will introduce the mathematical equations, the boundary conditions as well as a section on the numerical validation of the theoretical results obtained in [8] and in [9].

### 3.1 CSF models

In this section we introduce the lumped models for intracranial dynamics (see [21], [25]) for which we are going to perform numerical simulations. They are based on a mechanical description of the CSF flow and on the mechanical interaction between the parenchyma and the CSF compartments. In particular we consider the simplified formulations of the Linninger’s model ([21]) which have been analyzed in [8] and in [9].

The first model reads as follows

**Model A**

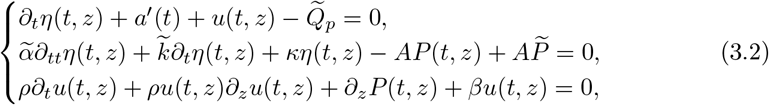

where 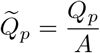 and 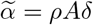, since the axial section, *A*, is considered constant in A this model.

Moreover, we want to treat numerically the following two models, obtained by improving Model **A** (see [9]).

**Model A1**

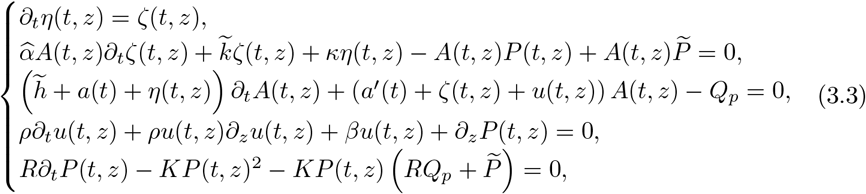

**Model A2**

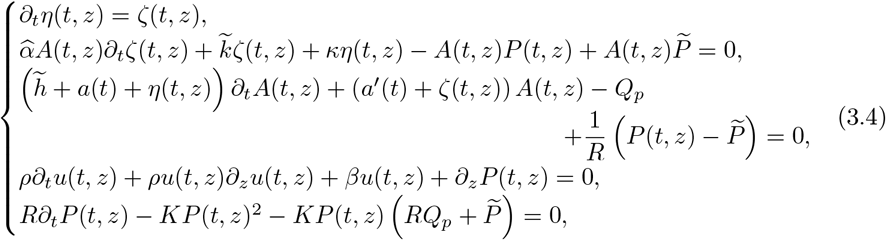

where 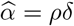 and the cross sectional area is not a constant.

In all the previous models the time *t* is such that *t* ∈ [0, *T*_0_] and *z* ∈ [0, *L*]. In particular, since the equation (3.3)_5_ shows a blow up at the finite time

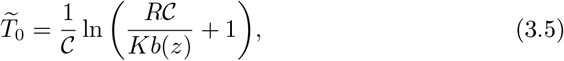

where 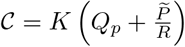, for the systems (3.3) and (3.5) we consider 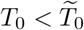.

Models **A, A1** and **A2** are lumped parameter models based on averaging axially the Euler equations with the addition of some simplifying assumptions, so they are essentially a physically based description. The overriding consideration in this context, is the complexity of the CSF fluid-structure interaction and the heterogeneity of the compartments perfused by the CSF, therefore they need to be studied by patching together several elementary tracts. This approach is based on classical continuity arguments and may be used for an accurate description of several segments of the CSF circulatory network.

In the aforementioned models the equations (3.2)_1_, (3.3)_3_ and (3.4)_3_ describe the continuity of the CSF flow in the ventricles, while (3.2)_3_ is the axial momentum equation along a streamline in the flow direction and with its nonlinearity represents the core of the theoretical analysis of the systems (3.2), (3.3) and (3.4). The equations for the acceleration of the elastic tissue are (3.2)_2_ and (3.3)_2_ and the evolution of the intracranial pressure is characterized by (3.3)_5_.

It is important to point out that the three models represent a first attempt of a pure mathematical analysis concerning system of equations which describe CSF fluid dynamics, therefore they are derived by assuming simplified properties and by neglecting particular interactions with the vascular system (for more details see [8] and [9]).

The following discussion of the numerical approach will establish the boundary conditions we will assume in order to perform proper simulations as well as the numerical feedback of the theoretical results obtained for Models **A**, **A1** and **A2**.

#### 3.1.1 Initial and boundary conditions

In order to be consistent with the notation adopted for the analysis of Models **A, A1** and **A2** in [8] and [9], we recall the following initial data, fixed for the theoretical analysis, which are also required for the numerical simulations.

For Model A they read as follows

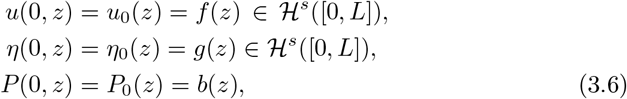

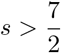, where

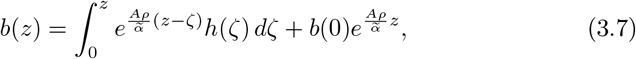

with

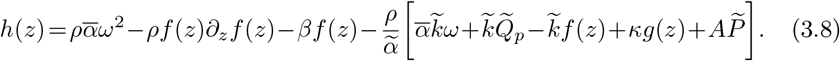

For Models **A1** and **A2**,

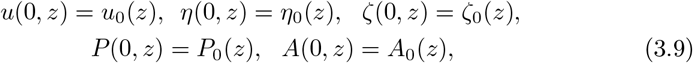

in 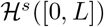 with 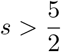.

The prescription of the data we need as boundary conditions for our 1*D* models is a very delicate task in the treatment of the physiological fluid flow problems. The measurement in vivo is the technique which is applied in order to collect clinical data. In general, measures may refer to pointwise velocities on a small portion of inflow/outflow boundaries of a fluid district or indirectly to average quantities like the flow rate over a cross section; for the pressure, data that are retrieved by means of noninvasive measures are almost invariably an average information over a section of interest. The most of the practical difficulties in the detection of the data occurs at the boundaries which characterize the outlet of the sections considered and unfortunately no information is available. These practical problems clearly add specific issues to the boundary treatment of mathematical and numerical models.

Models like Models **A, A1** and **A2** seem to be less difficult to treat even in absence of boundary data, indeed the mean value in space of a quantity in every point of the boundary makes the continuous problem well-posed. Nevertheless, the hyperbolic nature of the 1*D* model still raises issues of prescribing conditions at the outlets in a way consistent with the propagation dynamics. Moreover, the numerical discretization usually requires extra conditions which are not accounted in the continuous problem and that need to be assumed in a consistent fashion with the original model.

In Model **A** it has been assumed that the cross sectional area, A, is affected only negligibly by the pressure variation and represents for us a constant that we choose in the range of ~ 3 – 4 mm^2^ according to the clinical data and the section of the analyzed compartment. Furthermore, the boundary conditions involved in numerical simulations are selected by taking into account the following conditions (see Theorem 2.1.1 in [8])

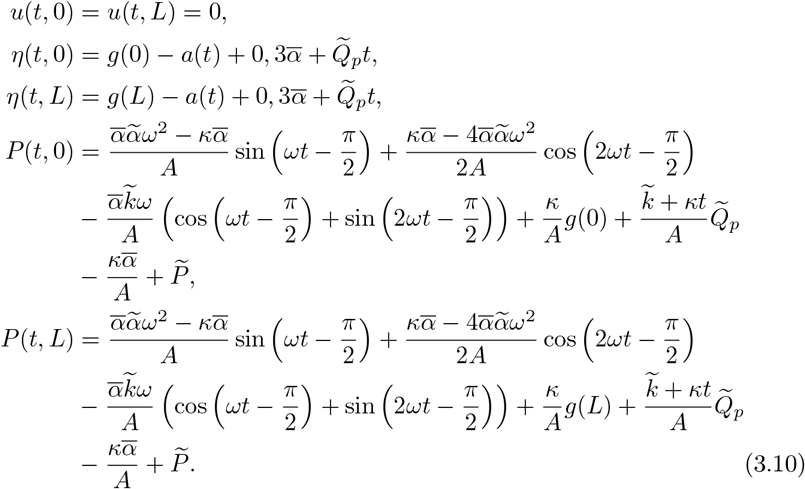

For Models **A1** and **A2**, we prescribe the boundary conditions

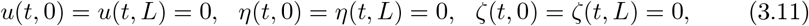

while, the boundary conditions for the pressure and the axial section are the following

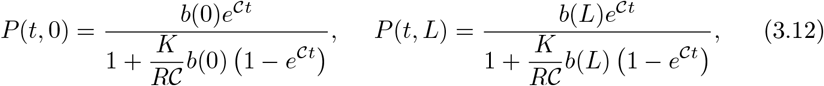

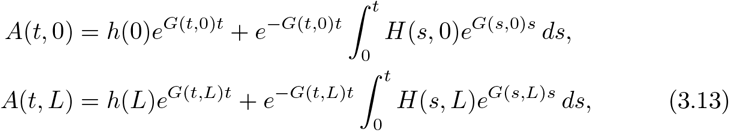

where

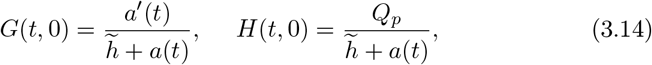

in Model **A1** and

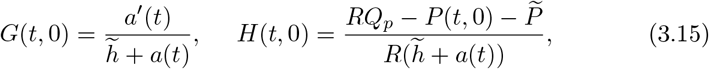

in Model **A2**.

The introduction of the no-slip conditions in the three models is clearly consistent with observations of macroscopic quantities such as the CSF flow rate through a circular capillary under a given pressure drop.

## 4. Goal of the numerical simulations

The aim of the present paper is to perform proper numerical simulations in order to complete and confirm the analysis provided in [8] and in [9], which represent a first attempt of studying cerebrospinal fluid models from a mathematical point of view. The theoretical results, whose reliability we want to prove with the aid of the simulations, concern the existence and uniqueness of the solutions for Models **A, A1** and **A2** that describe the dynamics of the CSF in the intracranial pattern by assuming suitable simplified assumptions that do not reflect totally the real physiology of the process examined. In detail, it has been proved that

- Model **A** admits a unique solution global in time if and only if

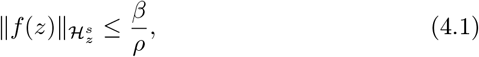

where *f*(*z*) = *u*(0, *z*) (for more details see [8]);
- there exists respectively a unique global solution to Models **A1** and **A2** provided that

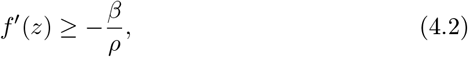

and

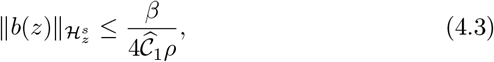

if *b*(*z*) < 0 with *b*(*z*) = *P*(0, *z*), and

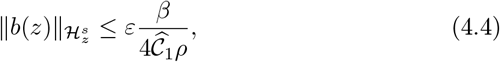

if *b*(*z*) > 0, where 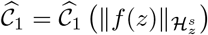 and 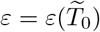 are constants (see [9]). Therefore, for all the models considered, we need proper smallness of the initial data in order to obtain global existence and uniqueness of solutions and this is the main result we want to prove also numerically in order to guarantee a more comprehensive overview since, as far as we know, there are no other outcomes in this direction in literature.

## 5. Numerical approximation

This section is focused on presenting the numerical results obtained by the models previously described. The first part of the section shows in detail the implementation of the problem in Matlab. The second part presents a variety of results dealing with the theoretical results obtained in [8] and [9]. Particular attention was devoted to analyzing pressures and flow patterns in the CSF compartments.

### 5.1 Numerical scheme for Model A

Let *D* = [0,1] be the computational domain, *a* and *b* constants such that 0 < *a* < *b* < 1, and Ω = [*a, b*]. Let *N* ≥ 1 be a fixed integer and *h* = 1/*N* the spatial step, let *D_h_* = {*x*_0_ < *x*_1_ < … < *x_N_* = 1} be the set of equally spaced grid points, and Ω_*h*_ = *D_h_* ∩ Ω the set of inside grid points.

Let *l* and *r* be such that *x_l_* ≤ *a* < *x*_*l*+1_, *x*_*r*-1_ < *b* ≤ *x_r_* (see Figure 5.1). We use a ghost-cell method to discretize the problem.

Let us begin to discretize (3.2) in Ω_*h*_ = {*x*_*l*+1_,…, *x*_*r*-1_}. We use a central difference scheme in space and forward Euler method in time for (3.2) obtaining:

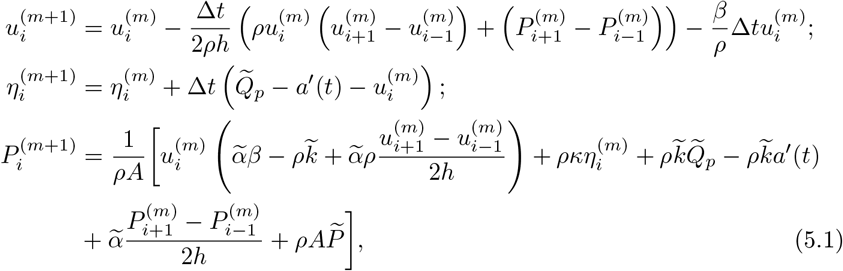

where *i* = *l* + 1,…*r* – 1 and all the non discretized terms are known values. Taking the maximum time step consented by CFL condition (see [5]), i.e. Δ*t* = *h*/2, we obtain in particular

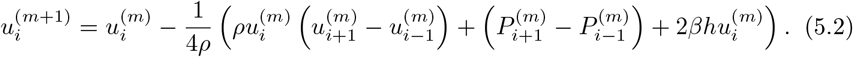

Furthermore, we choose a time step size of 5 × 10 ^−3^.

*Remark* 5.1. Explicit methods, like the forward Euler method, are very easy to implement, however, the drawback arises from the limitations on the time step size to ensure numerical stability ([29]).

*Conditional stability* concerning the existence of a critical time step dimension able to induce the occurrence of numerical instabilities, represents a typical property of explicit methods such as the forward Euler scheme. When explicit methods stability requires conditions too restrictive on the size of the time step, it’s recommended to adopt implicit methods. Nevertheless, in order to treat nonlinear problems implicit methods require an expensive computational cost since 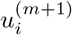 is defined only in terms of an implicit equation.

The model discretized numerically in (5.1) and (5.2) is characterized by a system of linear and nonlinear equations for which a more stable numerical method than the Euler’s one is recommended. But in this case we noticed that the latter method works perfectly such as the Matlab solver *ode45* since the intracranial pressure is able to dampen the behavior of the CSF flow velocity and plays the same purpose as a small time step size for the Euler method that guarantees the solutions remain inside the stability region.

#### 5.1.1 Numerical simulations

Based on the discretization set up in the previous section, in what follows we want to carry out the simulations for Model **A**.

For the initial data we proceed in the following way:

a. for the tissue displacement, *η*, we fix

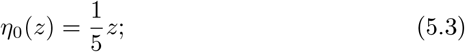
b. the pressure at the initial time, *P*_0_(*z*), is implemented exactly as required by the conditions (3.7) and (3.8);
c. for the axial CSF flow, *u*, we choose

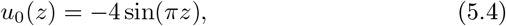

a negative initial datum which is in the “ safe zone” required by the global existence condition for Model **A** and that will show a different behavior with respect to the one performed in [8].

Therefore the previous initial conditions on pressure and flow velocity satisfy the global existence theorem stated in [8].

In Figure 3 we can observe in a section of length *L* = 1 the profiles of the pressure (red), the tissue displacement (green) and the flow velocity (blue) up to the final time *t* = 0.5.

**Figure 3:**
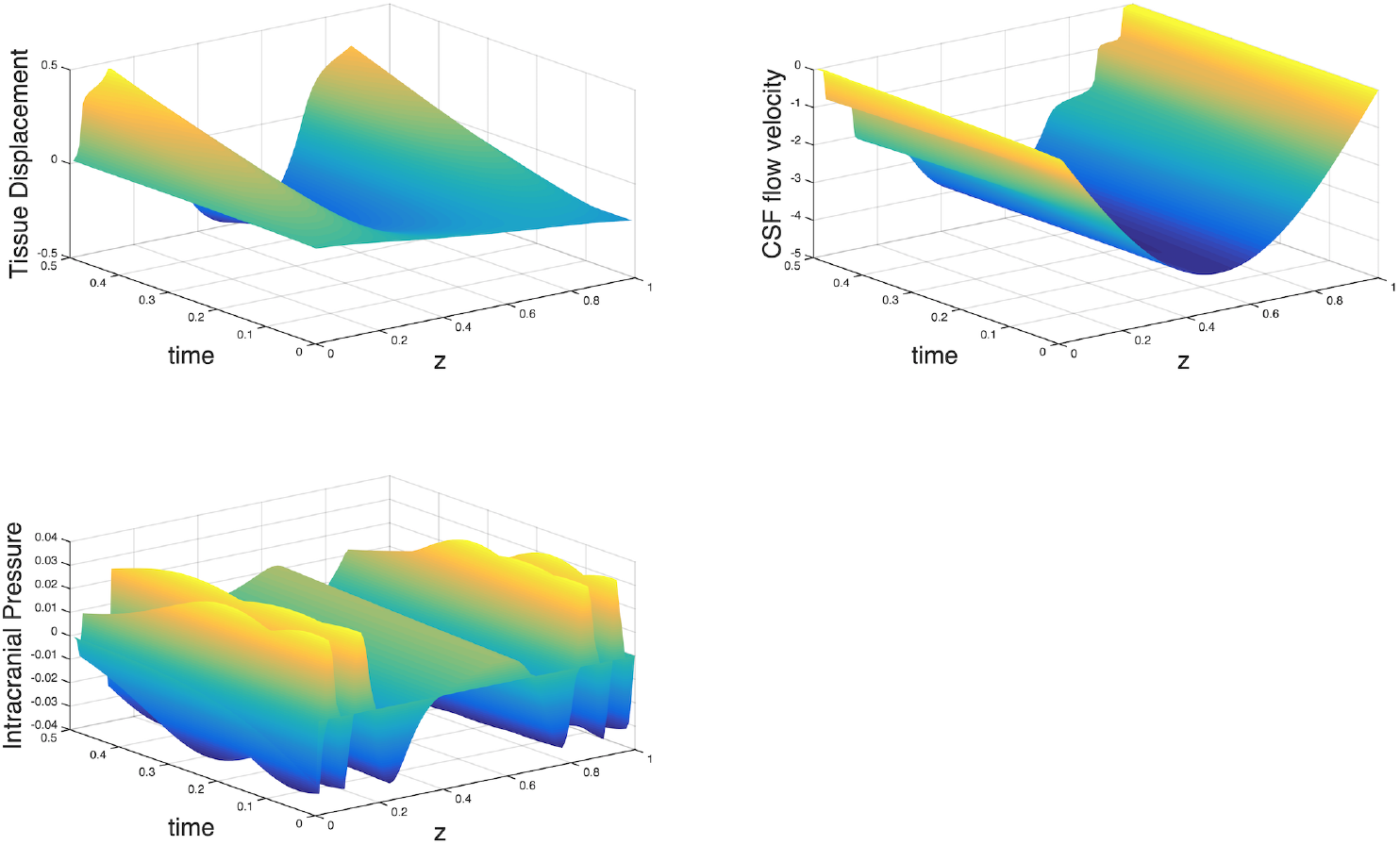
The evolution of *P, η* and *u*.

First of all, it’s very interesting to observe the behavior of the CSF flow velocity which shows an inverse fluid flow motion in the plot: this scenario is validated by the physiology of the CSF which is a pulsatile fluid affected by the heart beat and by the anatomy of the cerebral compartments that interact dynamically, therefore, in CSF circulation a small quantity of fluid is allowed to flow backwards inside the compartments (see [6], [7]). The tissue displacement evolves in a realistic fashion by means of the stress and the strains related to the fluid motion performed. The intracranial pressure, which is completely defined by the evolutions of the other quantities such as expressed in the system of equations (3.2), is represented by a profile very similar to the common signals detected in the in vivo measurements.

The second step consists in investigating the same framework described by Model **A** when the global existence condition is violated.

Therefore we fix the following initial datum

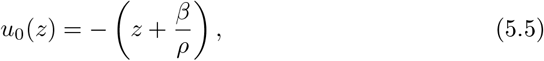

in order to observe what happens if we violate the global existence condition with a datum that is very close to it.

In Figure 4 the plots of the three quantities show a simultaneous blow up, a behavior that is in perfect agreement with respect to the study performed in [8].

**Figure 4:**
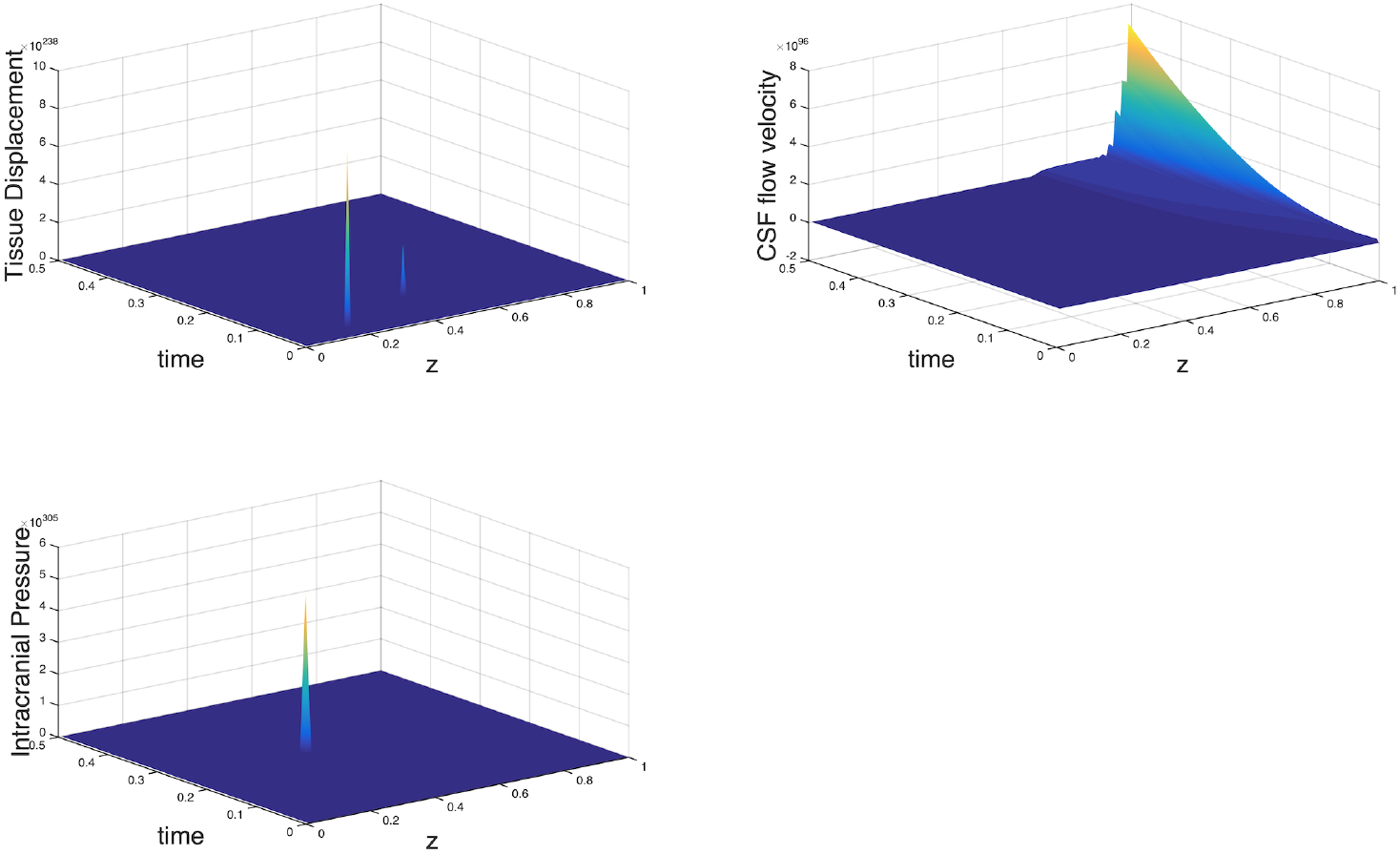
The blow up of the CSF flow velocity which occurs by violating the global existence condition.

Therefore, both simulations 3 and 4 support the conclusions achieved with the global existence theorem proved in [8].

### 5.2 Numerical approach for Models A1 and A2

This section is devoted to the numerical approximation of Models **A1** and **A2**. There exists, in this regard, a wide range of approaches and, among the many numerical methods useful to our aim, we decide to adopt the Runge-Kutta MATLAB solver *ode45*. This choice is due to the fact that we need to follow a different strategy with respect to the numerical schemes employed for Model **A**. The latter model, as explained in Section 5.1, does not require more stability in methods than the ones adopted in order to carry out the corresponding simulations, while in Models **A1** and **A2** we are considering a more comprehensive intracranial pattern, then more quantities as well as a further equation for the evolution of the intracranial pressure come into play in both models, which should be treated numerically by adopting suitable stable schemes as the solver *ode45*.

The function *ode45* implements 4th and 5th order Runge-Kutta formulas with a variable time step for efficient computation and is designed to handle the following general problem:

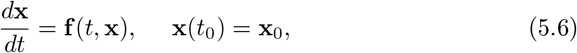

where *t* is the independent variable, x is a vector of dependent variables to be found and x(*t*_0_) is a function of *t* and x. The mathematical problem is specified when the vector of functions on the right-hand side of the equation (5.6), is set and the initial conditions, x = x_0_ at time *t*_0_ are given.

To this aim we define for both models a vector **X** as follows

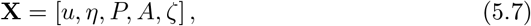

with

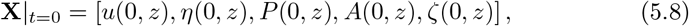

where *u, η, P, A, ζ* are defined themselves as vectors, as well. There is no preference for placing the variables, in fact, any arbitrary order is perfectly valid.

We adopt the same space and time discretization defined in the previous section and we create two extra ghost cells, one just at the left of 0 and one just at the right of 1. This procedure is used for convenience, since it simplifies the implementation of boundary conditions a lot.

At this point, the systems (3.3) and (3.4) can be easily written as systems of first-order differential equations in the following way

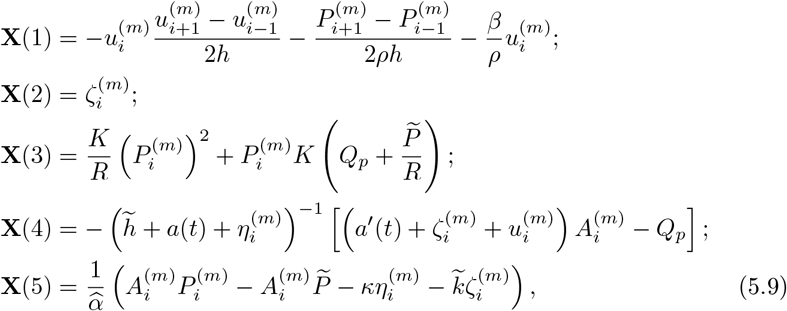

for Model **A1** and

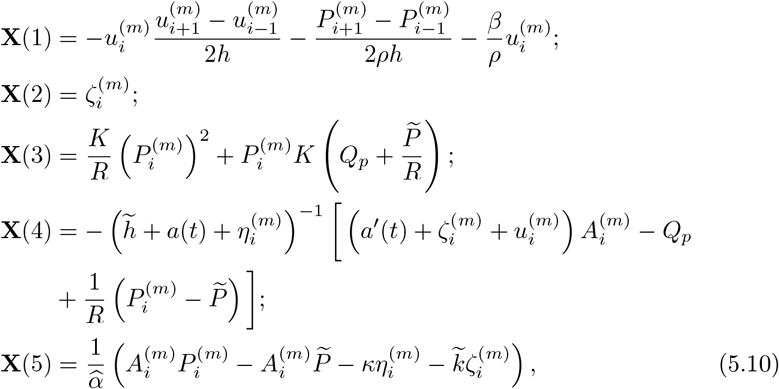

for Model **A2**, where *i* = *l* +1, … *r* − 1 and all the non discretized terms are known values.

Of course, the notation **X**(*j*) with *j* = 1, … 5, denotes an entire vector, not only a single value, in particular *length*,(**X**(*j*)) is equal to the length of every variable vector (*u,η, P, A,ζ*) respectively, in other words each column of **X** is a different dependent variable.

The routine *ode45* integrates the systems of ordinary differential equations (5.9) and (5.10) over the interval *T*_0_ =0 to *T_final_* = 1, with initial conditions (5.8), it uses a default tolerance and displays status while the integration proceeds.

In our simulations the initial mesh size is determined by *dz* = *L*/*nz*, where, for the sake of simplicity, *L* is fixed equal to 1 and nz is the total number of meshes that we take equal to 200. Finally we set the time discretization with Δ*t* = 5 × 10^−3^.

#### 5.2.1 Numerical simulations

In order to run numerical simulations for Models **A1** and **A2**, we assume

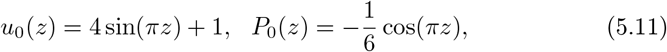

which are not so far from a good real approximation of the CSF flow velocity in a small compartment perfused by the fluid. In particular, we adopt a negative initial datum for the intracranial pressure in order to distinguish the present simulations to the ones already carried out in [9].

As a byproduct of our numerical results, we obtain interesting informations about the behaviors of the quantities involved in these models.

As a second task, we implement again the models numerically but we fix initial data *u*_0_(*z*) and *P*_0_(*z*) which violate simultaneously the conditions of the global existence theorem ([9]). Furthermore, we analyze the behavior of the models when the condition of the pressure solely, (4.3), is not satisfied while the costraint on the flow velocity, (4.2), is fulfilled and vice versa. We shall compare these simulations to the analysis developed in [9] and show the accuracy of our approach.

**First case**: the initial data fulfill the global existence conditions (4.2) and (4.4).

We perform the simulation by assuming the initial condition (5.11) and arbitrary data for the others quantities for which we choose

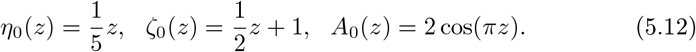

In Figures 5 and 6 we can observe the parabolic profile of the CSF flow velocity of both models due to the no-slip boundary conditions assumed. This quantity shows a slow variation in relation to the fluid motion.

**Figure 5:**
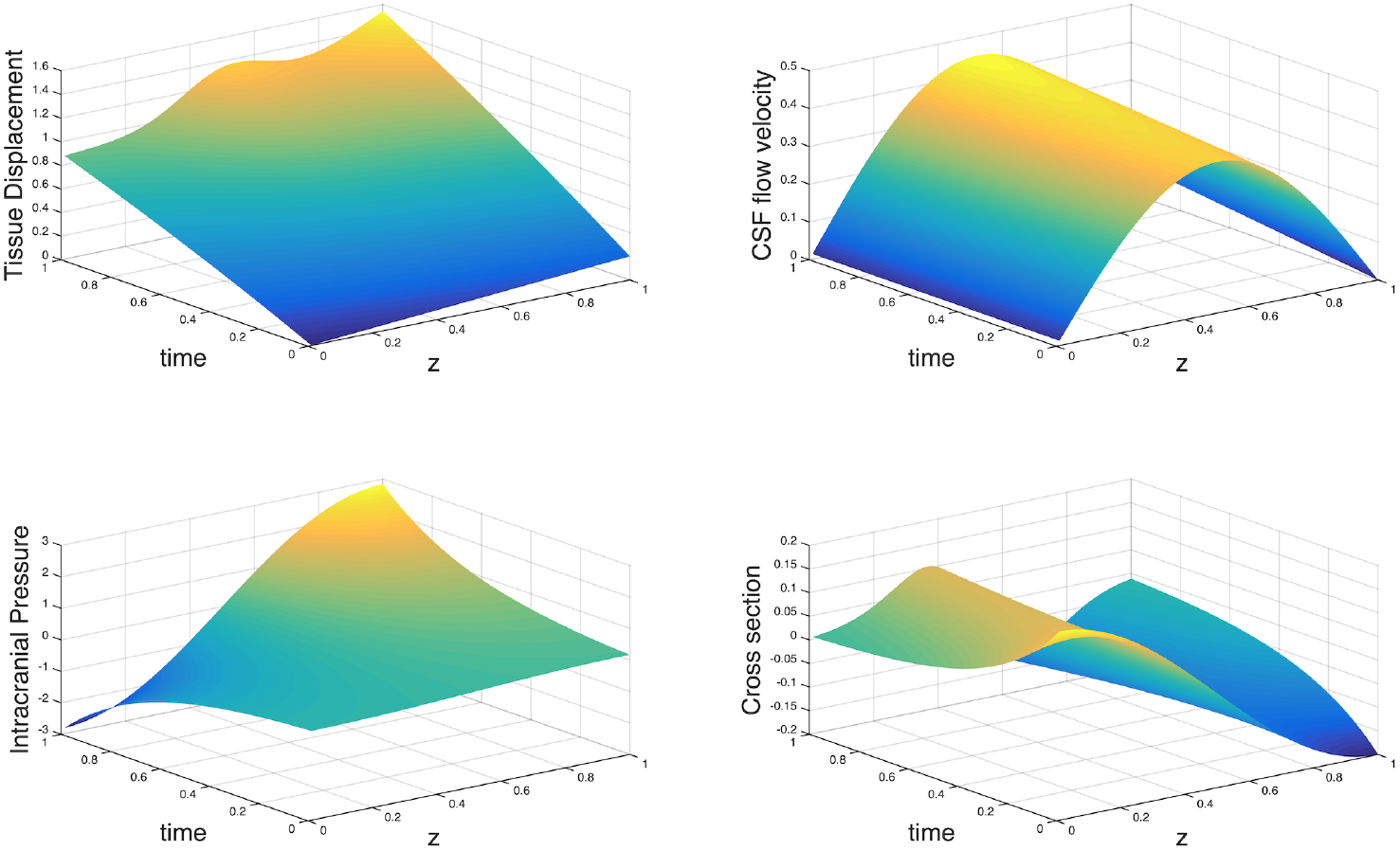
Evolution of *P, η, u* and *A* for Model **A1**.

**Figure 6:**
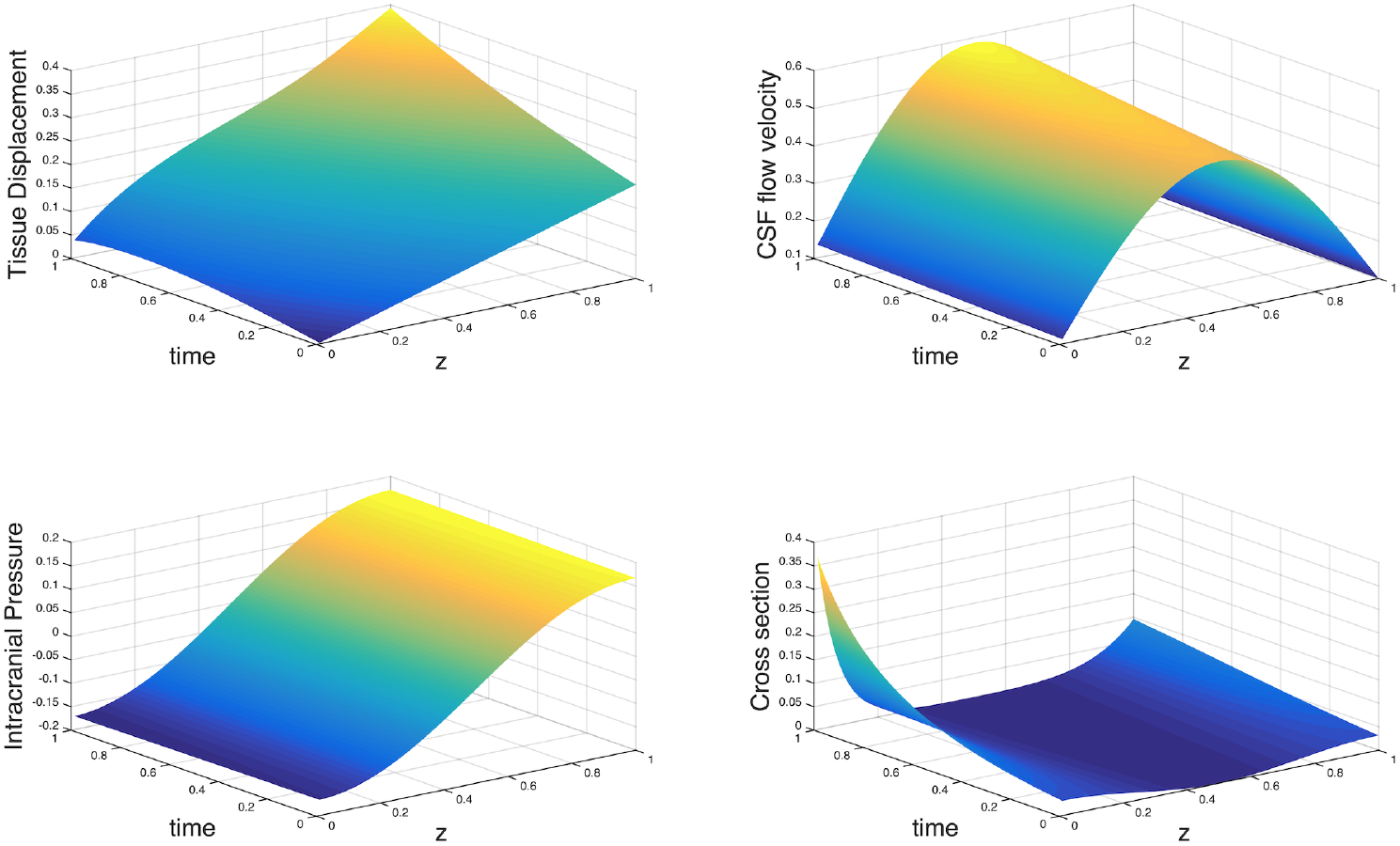
Evolution of *P, η, u* and *A* for Model **A2**.

The plot of the tissue displacement, *η*, highlights small contractions and dilatations of the tissue (see Fig. 5) in Model **A1**, while in Model **A2** the displacement displays a remarkable increase in the real physiological values of tissue displacements (see Fig. 6). For A we can observe a drastic reduction in the cross sectional area in Model **A2**, while in Model **A1** the axial section increases rapidly in the first part of the section until a first maximum point, then there’s a drop and a small increase towards the boundary. This behavior is a consequence of the negative intracranial pressure we fixed as initial datum. Indeed the intracranial pressure, which is affected by small variations in Model **A2** and touches big numerical values in Model **A1**, acts a compression in the cerebral pattern and the other quantities involved in the models show a physiological reaction with respect to the activity of *P*.

Therefore the parameters behave as predicted by the analysis performed in [9] even if the assumptions on the boundary conditions affected the simulations that appear clearly different from the real physiology of the CSF flow.

**Second case**: the initial data violate the global existence conditions (4.2) and (4.4).

I. **Violation of both global existence conditions**. We assume

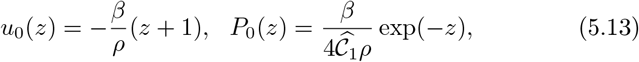

where 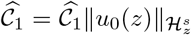 is the constant that appear in the global existence theorem proved in [9] (see Theorem 2.5). The previous initial conditions are assumed in order to show which is the behaviors of the quantities involved in both models when we fix the data at time *t* = 0 exactly equal to the global existence conditions required by the mentioned theorem. We have no costraints on the initial data of *η* and *A*. Therefore, henceforth, we adopt the conditions (5.12) for the other variables of the models since they are close to the experimental data and describe properly the initial framework of the CSF dynamics in a single compartment. In Figure 7 we can observe that the blow up has been achieved after several time steps but the shock waves appear simultaneously with significant numerical values. While, Figure 8 shows that in Model **A2** the different quantities blow up after only five time steps and this means that the initial conditions (5.13) affect more strongly the behavior of the model since the intracranial pressure interact also in the form of the reabsorption term.
II. **Violation of the global existence pressure condition**. As initial conditions we fix

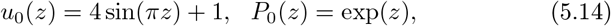

for the flow velocity and the pressure. In Fig. 9 it is possible to observe that the pressure blow-up leads to a simultaneous blow-up of the other quantities involved in the simulation: this means that such pressure is able to amplify the motion of the cerebrospinal fluid throughout the small compartments and thus to affect significantly the entire intracranial fluid dynamics. Fig. 10 shows a drastic pressure drop in Model **A2**: even though the input pressure refers to a recumbent position, it blows up by taking negative values. We can notice that

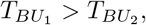

where *T*_*BU*_1__ is the blow-up time of Model **A1** and *T*_*BU*_2__ is the blow-up time of Model **A2**; this is due to the fact that the reabsorption rate, *Q_a_*, is affected by the pressure and accelerates the blow-up process in Model **A2**.
III. **Violation of the global existence flow velocity condition**. For the last simulations we assume

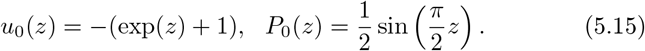 Figs. 11 and 12 show that the blow-up is achieved after almost the same number of time steps, but in Model **A2** it is possible to observe that the pressure is more affected by the violation of the global condition of the flow velocity. The reason can be found, even then, in the particular physiology described by Model **A2**: the CSF reabsorption requires the employment of a comprehensive set of cerebral mechanisms in which the pressure plays a fundamental role. Even though we are not violating directly the condition on the ICP, the reaction of the pressure is immediate since the latter is strictly connected to the flow velocity, and its behavior in these simulations is consequence of the relation between the global condition (4.3) and the flow velocity condition (4.2) due to the constant 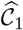 that appears in the former condition.

**Figure 7:**
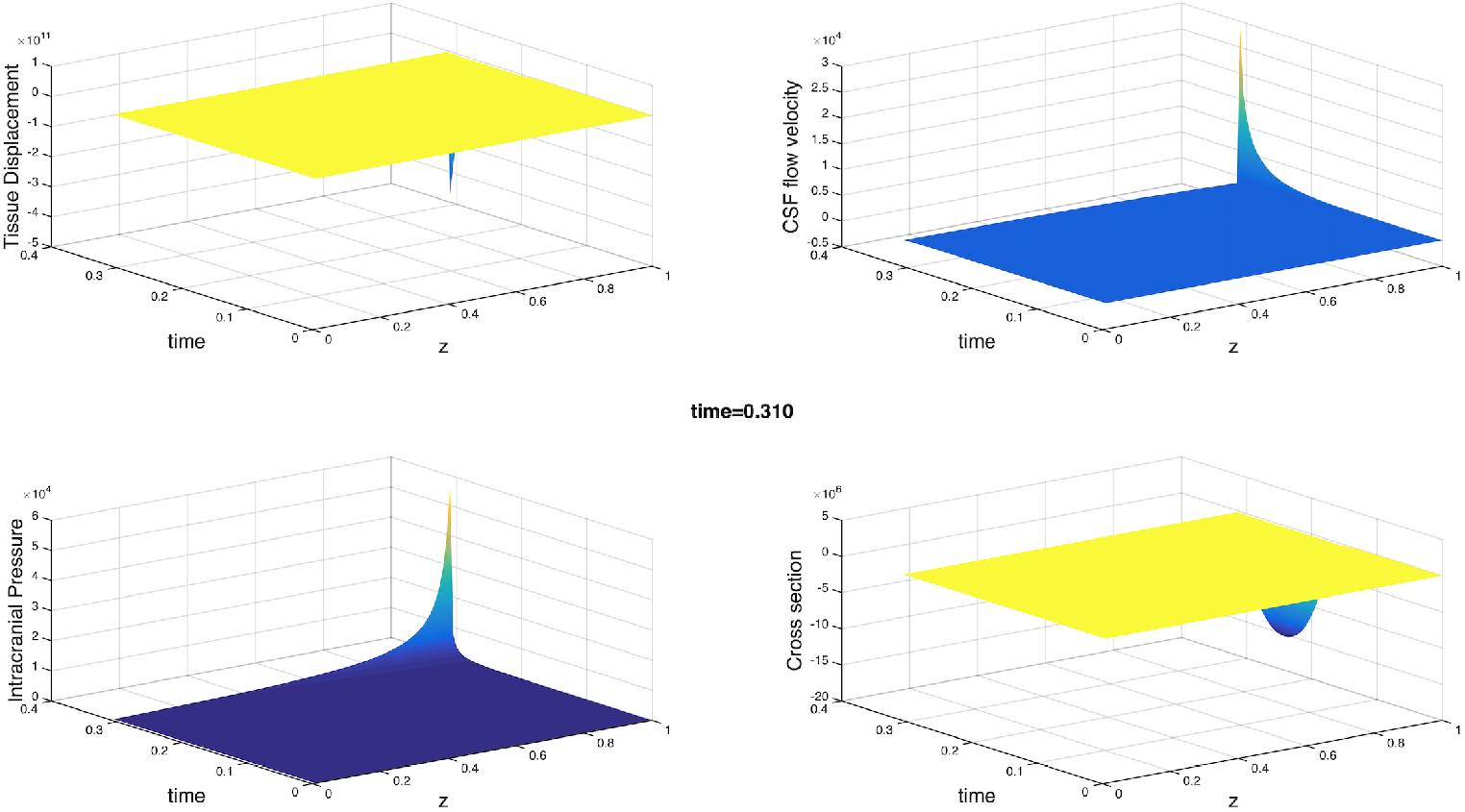
Blow-up which occurs in Model **A1** with initial conditions (5.13), (5.12).

**Figure 8:**
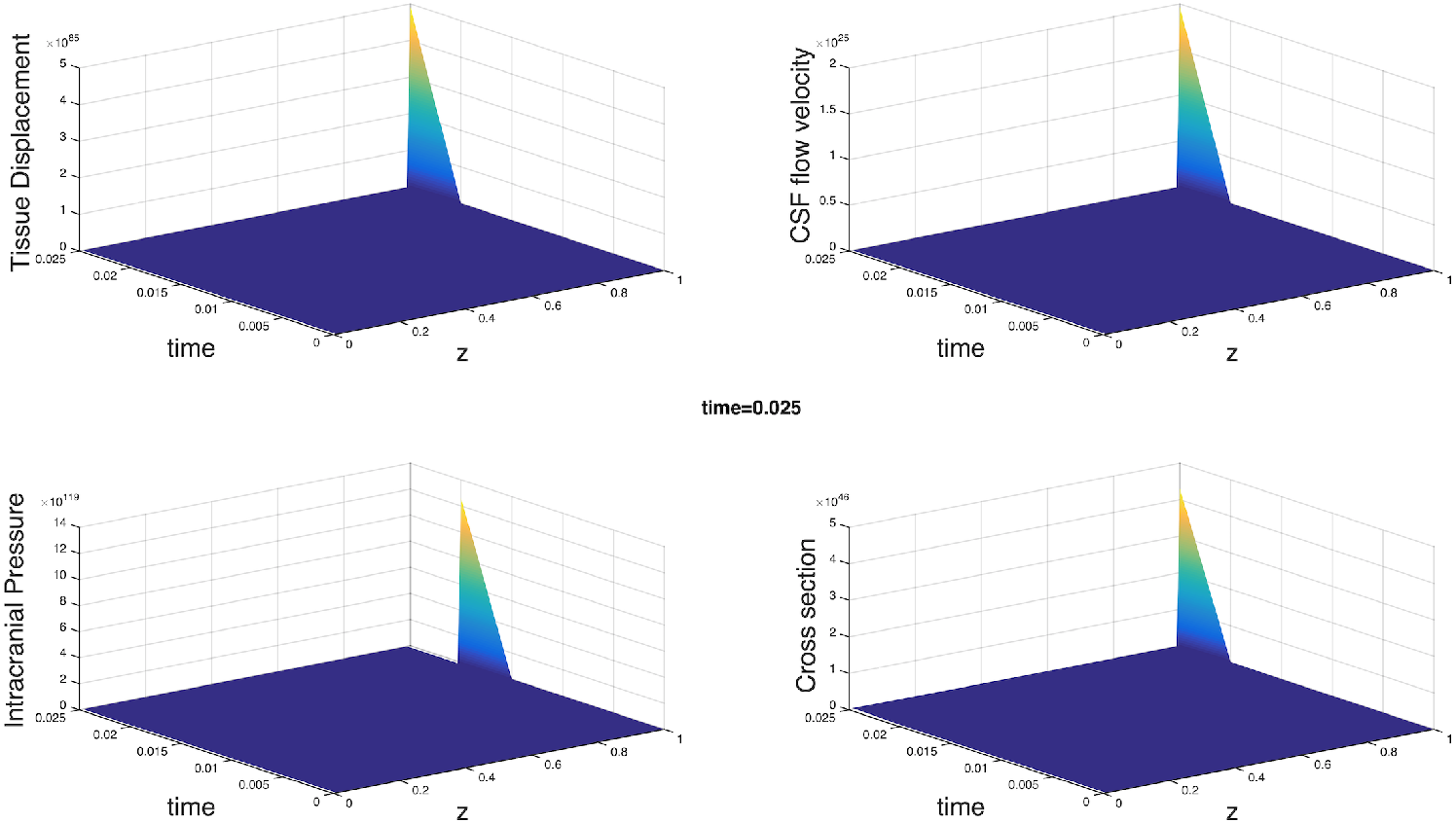
Blow-up which occurs in Model **A2** with initial conditions (5.13), (5.12).

**Figure 9:**
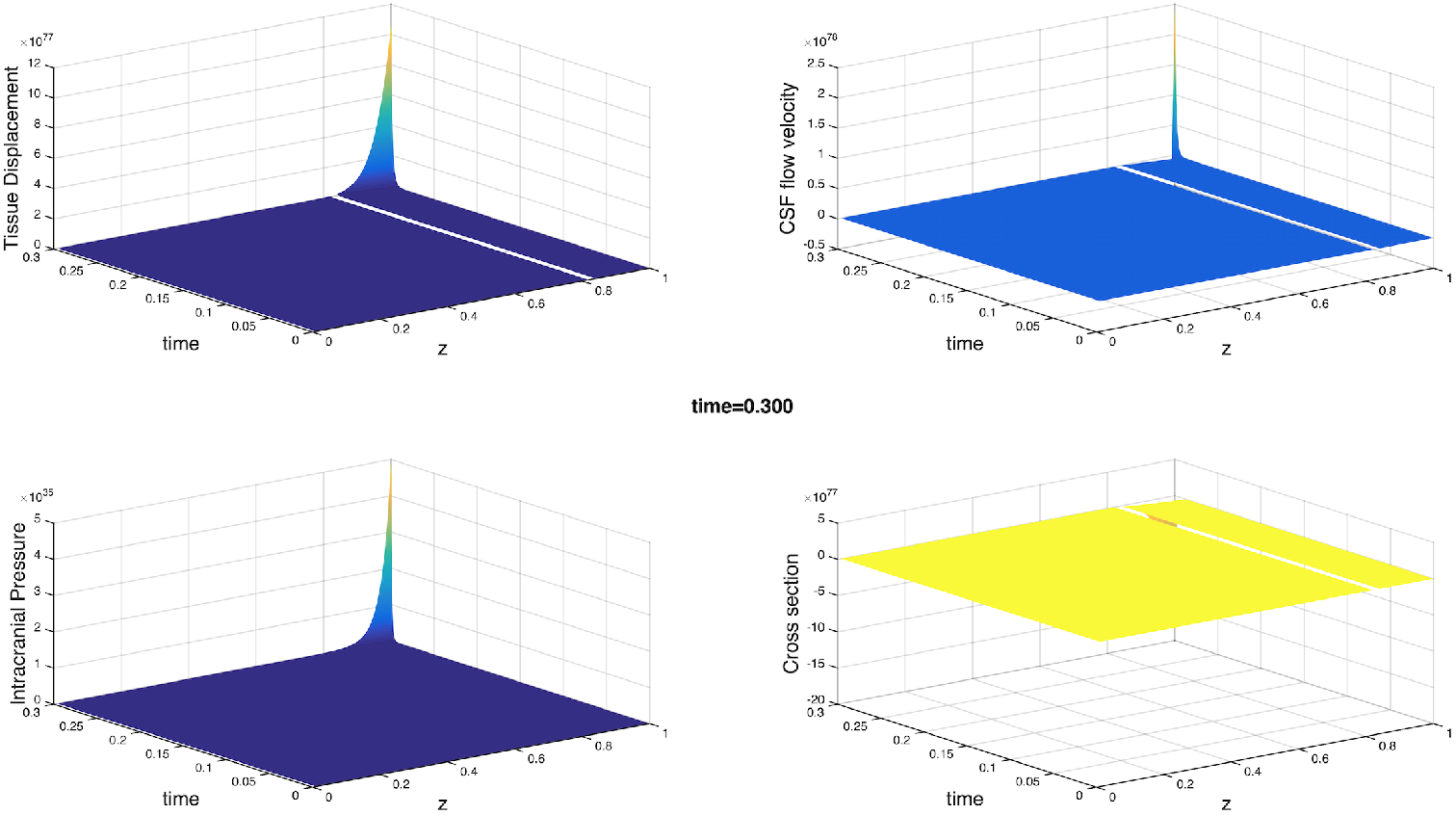
Blow-up which occurs in Model **A1** by violating the pressure condition (4.3).

**Figure 10:**
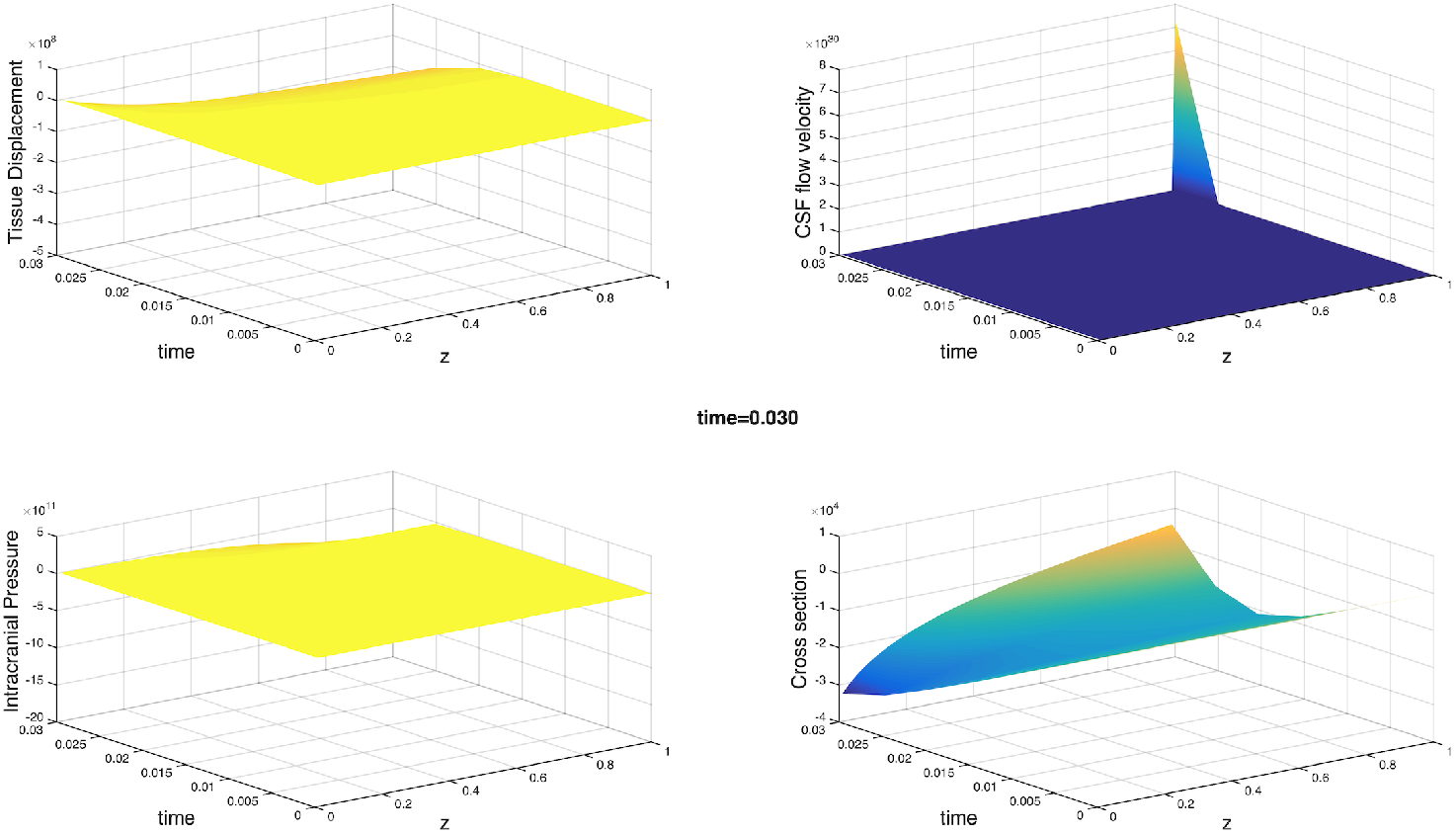
Blow-up which occurs in Model **A2** by violating the pressure condition (4.3).

**Figure 11:**
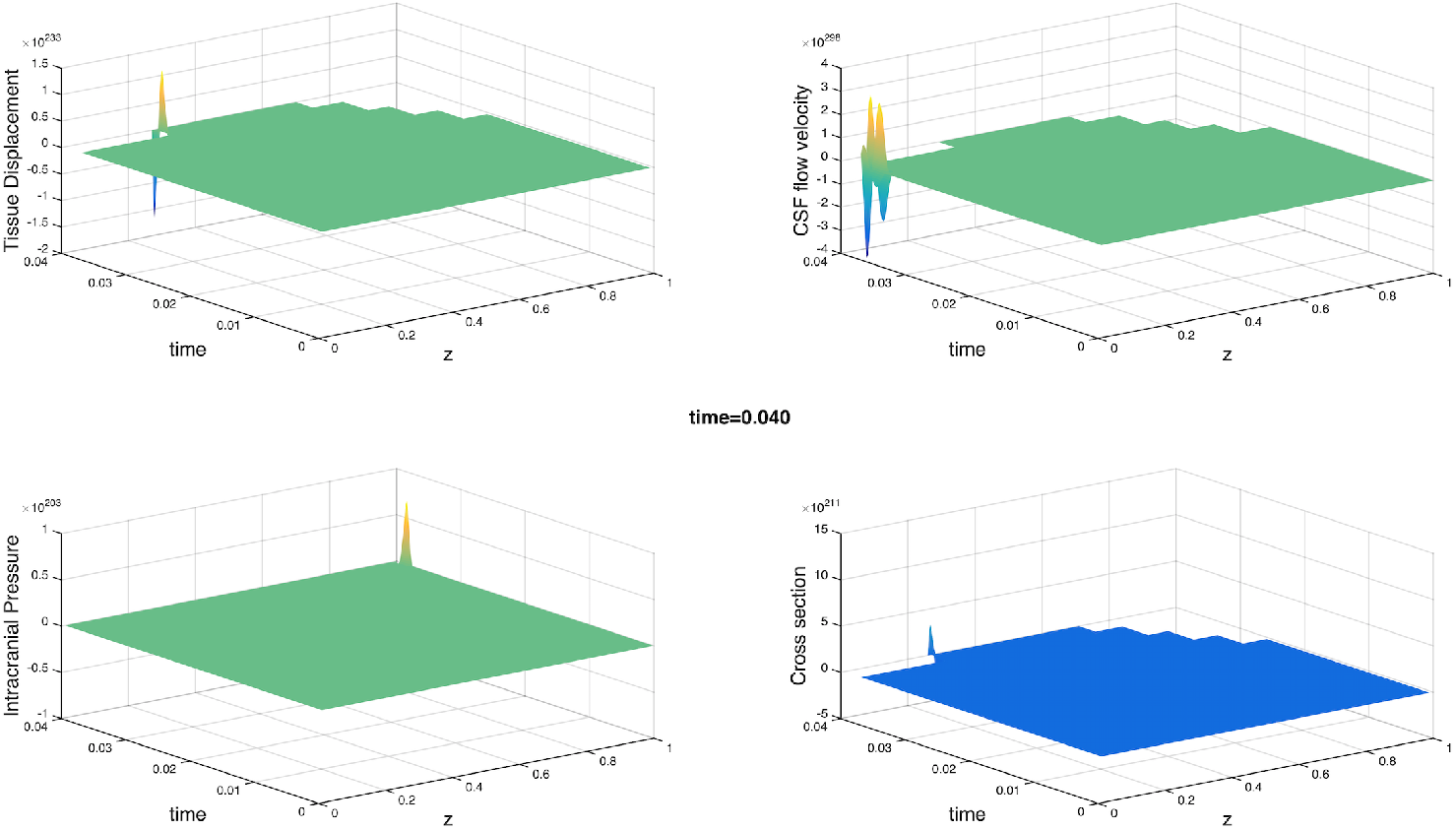
Blow-up which occurs in Model **A1** by violating the flow velocity condition (4.2).

**Figure 12:**
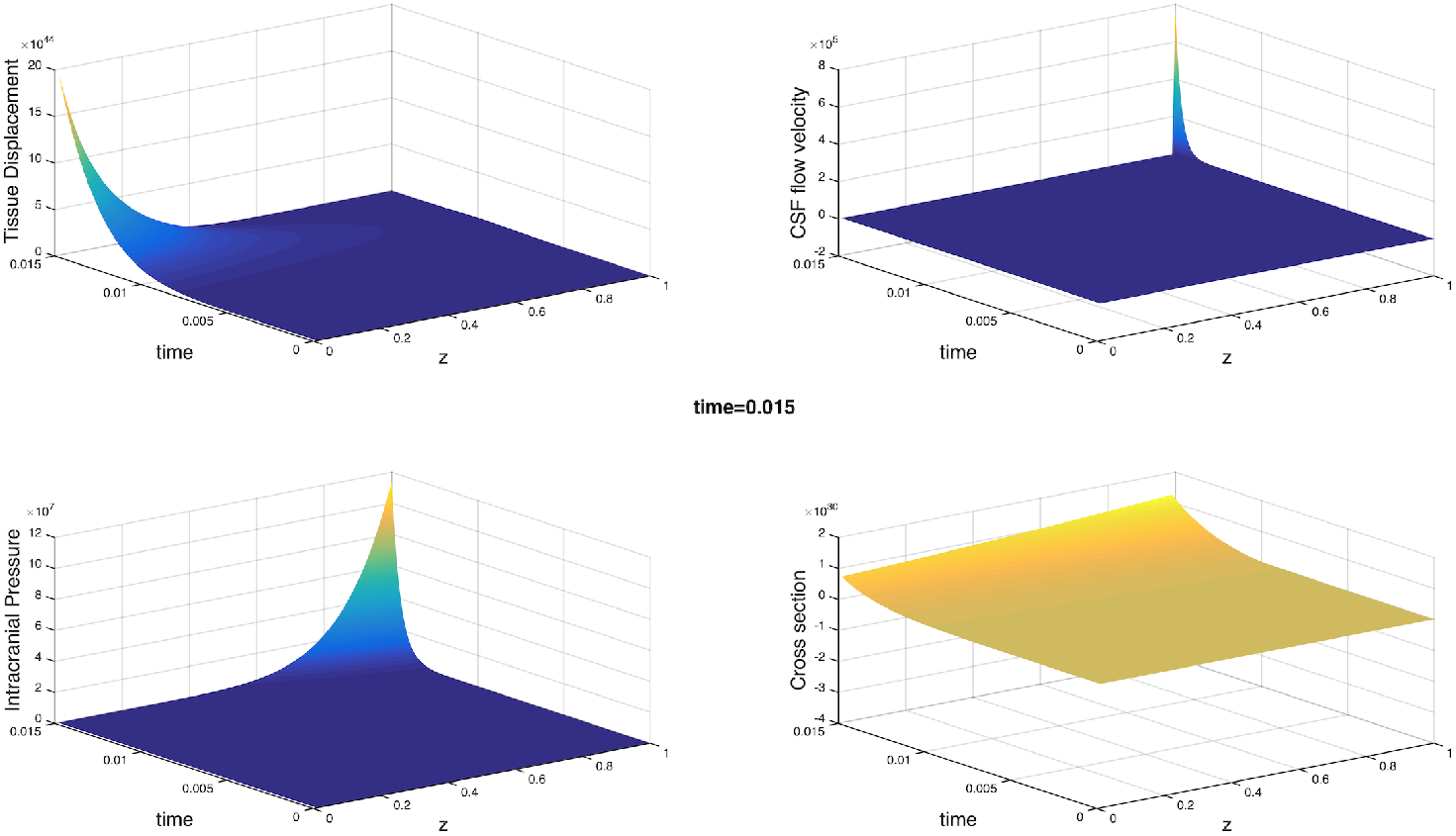

## 6. Final conclusions on numerical simulations

This paper has been designed to provide a first comparison of the CSF dynamic Models **A, A1** and **A2**, and a comprehensive overview of numerical simulations in order to prove the reliability of the mathematical analysis related to these models developed in [8] and [9]. The simulations have been carried out by adopting simplified assumptions on the boundary conditions and initial data consistent, as far as possible, with the experimental ones.

Two different numerical approaches for Models **A, A1** and **A2** are presented in order to realize accurate simulations and to avoid expensive computational costs. In conclusion, we can say that the performed numerical simulations support the theoretical results achieved in [8] and [9], and represent a first important step towards a new line of research in which fluid dynamics and mechanical models for the brain and its many components can be deeply investigated, not only from a purely empirical view point but also from a mathematical standpoint, in order to close the gap among different scientific fields and to guarantee an improvement on the research outcomes.

